# Neuronal glycolytic reprogramming drives lethality via accelerated aging in a *Drosophila* model of tauopathy

**DOI:** 10.1101/2025.10.27.684860

**Authors:** Richa Gupta, Hope McGinnis, Elham Rastegari, Matthew S. Price, Kartik Venkatachalam

**Affiliations:** Department of Integrative Biology and Pharmacology, McGovern Medical School at the University of Texas Health Sciences Center (UTHealth), Houston, TX, USA; Neuroscience Graduate Program, The University of Texas MD Anderson Cancer Center UTHealth Houston Graduate School of Biomedical Sciences; Molecular and Translational Biology Graduate Program, The University of Texas MD Anderson Cancer Center UTHealth Houston Graduate School of Biomedical Sciences

## Abstract

Neurometabolic dysfunction is a hallmark of Alzheimer’s disease (AD) and tauopathies. Whether these changes drive pathology or represent compensatory, protective responses remains unresolved. Here, we demonstrate that human tau induces Warburg-like metabolism in *Drosophila* neurons, characterized by coordinated upregulation of glycolytic enzymes and lactate dehydrogenase that mirrors metabolic signatures in human AD. Despite intact mitochondrial oxidative phosphorylation, *tau*-expressing fly neurons preferentially utilize glycolysis for ATP production and operate with diminished metabolic reserve. Crucially, this metabolic reprogramming drives rather than protects against pathology as genetic suppression of glycolysis or lactate dehydrogenase completely rescued tau-induced lethality. Further, Gompertz mortality analysis revealed that hyperactive glycolysis in tau neurons drives premature lethality by accelerating biological aging rate without affecting baseline mortality. Collectively, these findings establish aberrant neuronal glycolysis as a cause rather than a consequence of tau pathology, and demonstrate that sustained glycolytic metabolism in mature neurons exacts a specific cost in the form of accelerated aging.

## INTRODUCTION

Alzheimer’s disease (AD), tauopathies and other related causes of dementia are characterized by progressive neuronal dysfunction, cognitive decline, and increased mortality (1–3). While efforts at mitigating the aggregation of hyperphosphorylated tau and amyloid-β have dominated therapy development, brain metabolic dysfunction has emerged as an equally prominent feature of dementias (4–6). Yet the causal relationship between metabolic alterations and disease progression remains unresolved—*does metabolic dysfunction drive pathology, or does neuronal damage cause secondary metabolic failure?* Although recent studies of human tissue and patient-derived neurons have revealed transcriptomic and proteomic changes in AD neurons that are expected to augment glycolysis and promote lactate production (5, 7–9), the functional significance of these changes remains ambiguous. Glycolytic upregulation could represent either a protective compensation needed for ATP homeostasis or a maladaptive response that exacerbates pathology. Either interpretation has precedent. Given that neurons respond to diminished mitochondrial ATP production by upregulating glycolysis (10), increased glycolysis in AD neurons could be protecting against the bioenergetic deficits stemming from mitochondrial dysfunction (11). Arguing against interpretation, however, is the observation that healthy mature neurons actively repress glycolysis, instead utilizing glia-derived metabolites such as lactate and ketone bodies for fueling mitochondrial ATP production (12). In this view, ectopic induction of glycolysis in mature neurons might be detrimental to long-term neuronal viability.

We examined these scenarios using an established *Drosophila* model of tauopathies. Our previous work established expression of the 0N3R isoform of human *tau* in *Drosophila* glutamatergic neurons results in compromised ER–mitochondria Ca^2+^ transfer, a compensatory increase in IP_3_ receptor (IP_3_R)-mediated ER Ca^2+^ release that is maladaptive and underlies the shorter lifespan observed in those flies (13). Here, we combined transcriptomics, single-neuron and brain metabolic profiling, genetic manipulation, and quantitative survival analysis to further investigate the mechanisms of tau^0N3R^-induced pathology and mortality. We find that the transcriptional response associated with *tau^0N3R^* expression is reminiscent of that observed in human AD. Tau^0N3R^ drove coordinated upregulation of glycolytic enzymes and lactate dehydrogenase at transcriptional and protein levels, producing a Warburg-like metabolic shift that maintains ATP homeostasis through increased glycolysis and reduced reliance on oxidative phosphorylation (OXPHOS). Critically, we demonstrate that this metabolic reprogramming is not merely correlative, but rather causes premature mortality in flies expressing *tau^0N3R^* in neurons. Genetic knockdown of glycolytic genes or lactate dehydrogenase specifically in *tau^0N3R^*-expressing neurons restored normal lifespan. Additionally, using Gompertz mortality modeling to decompose survival into baseline mortality and aging rate parameters as reported in prior analyses of *Drosophila* longevity (14–16), we demonstrate that tau^0N3R^ accelerates biological aging without impacting baseline mortality. Importantly, we show that the accelerated aging rate stems from hyperactive glycolysis in neurons expressing *tau^0N3R^*. These findings establish neuronal metabolic reprogramming as a driver of tau neuropathology and reveal that sustained activation of glycolysis in mature neurons can exact long-term costs in the form of accelerated aging.

## RESULTS

### Transcriptomic analyses of *tau^0N3R^*-expressing *Drosophila* brains

In prior studies, we showed that neuronal expression of human *tau^0N3R^* in *Drosophila* results in significantly shorter adult lifespan stemming from cell autonomous increases in IP_3_R activity and ER Ca^2+^ dyshomeostasis (13). However, the molecular cascades initiated by tau^0N3R^ that culminate in organismal death remain incompletely understood. Given that tau^0N3R^-induced lethality manifests progressively during adulthood (13), we hypothesized that transcriptional changes preceding this phenotype would reveal initiating pathogenic events rather than the secondary consequences of neurodegeneration. To identify these early molecular alterations, we performed bulk RNA-sequencing (RNA-seq) on brains isolated from 3^rd^ instar larvae expressing *UAS-tau^0N3R^* (17) under the control of the ubiquitous *tubulin-GAL4* (Supplemental Tables 1A-1B). We used a ubiquitous rather than neuron-specific driver to maximize signal detection in whole-brain tissue because cell-type-specific expression would result in dilution of tau^0N3R^-associated transcriptional changes with those originating from cellular populations not expressing the transgene. Principal component analysis (PCA) of normalized counts for all 23,932 annotated *Drosophila* genes (Supplemental Table 1B) revealed striking separation of the transcriptional profiles of tau^0N3R^ brains relative to controls (Figure 1A). Strong divergence along PC1 indicated that tau^0N3R^ induced broad and fundamental alterations in gene expression patterns.

**Figure 1.**
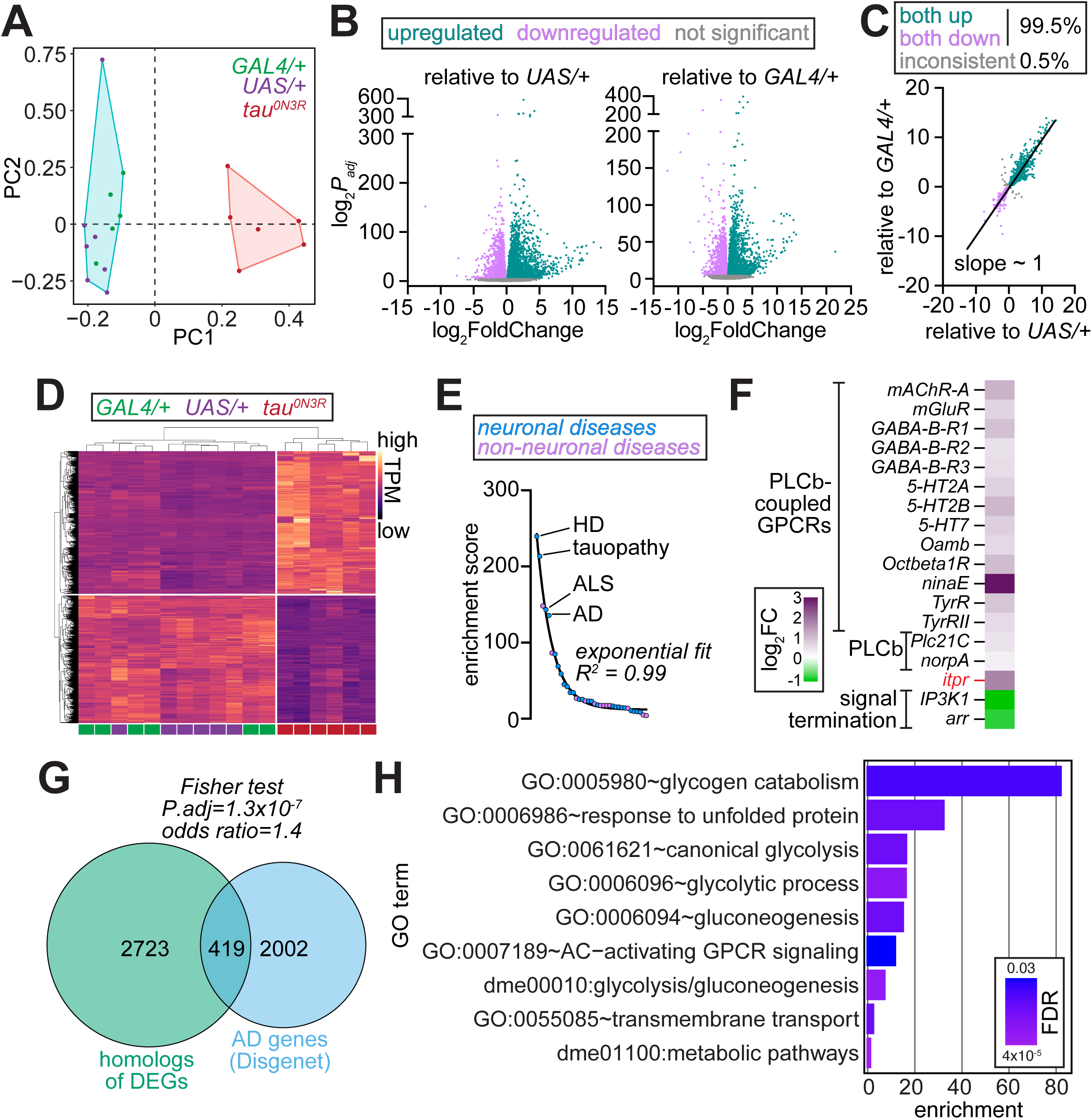
Transcriptomic analysis *tau^0N3R^*-expressing *Drosophila* brains reveals enrichment for pathways related to neurodegeneration and glycolytic upregulation. **(A)** PCA of normalized gene expression (TPM) from larval brains from *tubulin>tau^0N3R^* (red) and parental controls (*tubulin-GAL4*/+, green; *UAS-tau^0N3R^*/+, purple). Specification of two clusters in k-means clustering revealed separation of *tau^0N3R^*-expressing (red) and control (cyan) transcriptomes across PC1. **(B)** Volcano plots showing DEGs in *tubulin>tau^0N3R^* brains relative to *UAS-tau^0N3R^*/+ (left) and *tubulin-GAL4*/+ (right). Upregulated genes (teal), downregulated genes (magenta), and non-significant genes (adjusted *P*-value ≥ 0.05, gray) are indicated. **(C)** Scatter plot comparing log_2_ fold changes of DEGs identified relative to each control line. High concordance (slope ∼ 1) and 99.5% directional consistency demonstrate robust, genotype-specific transcriptional changes. **(D)** Heatmap of 4,519 high-confidence DEGs showing distinct clustering of tau^0N3R^ samples (magenta bars) from controls (purple and green bars). Color intensity in heatmap represents transcript abundance (TPM, normalized by row). **(E)** Enrichment analysis of tau^0N3R^ DEGs among genetic modifiers of *Drosophila* models of human diseases. Diseases are ranked by their enrichment score (odds ratio × −log_10_ adjusted *P*-value). Four of five are neurodegenerative disorders with enrichment scores dropping exponentially (R^2^ = 0.99, fit to exponential decay curve). **(F)** Expression changes of genes encoding PLCβ-coupled GPCRs, PLCβ paralogs, IP_3_R (*itpr*), and signal termination components (IP_3_ kinase-1 and β-arrestin). Heatmap shows log_2_ fold changes; upregulated genes (magenta) include signaling activators, while downregulated genes (green) include terminators. **(G)** Venn diagram showing overlap between human orthologs of tau^0N3R^ DEGs (green, 3,142 genes) and genes associated with Alzheimer’s disease from DisGeNET database (blue, 2,421 genes). Overlap of 419 genes represents ∼17% of AD-associated genes. Fisher’s exact test shows significant overlap. **(H)** Gene Ontology enrichment analysis of upregulated DEGs shared between brains of animals expressing *tau^0N3R^* and those overexpressing *itpr*. Bar length represents fold enrichment; color intensity indicates FDR.

Differential expression analysis identified 5,843 differentially expressed genes (DEGs) in tau^0N3R^ brains relative to *UAS-tau^0N3R^*/+ (3,145 up and 2,698 down), and 5,527 DEGs relative to *tubulin-GAL4*/+ (2,934 up and 2,593 down) (Figure 1B, Supplemental Tables 2A-2B). To distinguish tau^0N3R^-associated effects from those stemming from idiosyncratic background variations specific to each control, we intersected both sets of DEGs and identified 4,543 shared DEGs. 4,519 of these genes (∼99.5%) showed concordant directionality of gene expression changes in both DEG sets (Figure 1C and Supplemental Table 2C). This final high-confidence set, which we used for all subsequent analyses, was comprised of 2,385 upregulated and 2,134 downregulated genes (Figure 1D and Supplemental Table 2C).

### Tau^0N3R^-induced transcriptional changes reflect signatures of neurodegeneration and Alzheimer’s disease

To determine whether tau^0N3R^-induced transcriptional changes reflect broader neurodegenerative processes, we compared tau^0N3R^ DEG set to genetic modifiers previously identified in *Drosophila* disease models. FlyBase catalogs modifiers for 208 distinct classes spanning 283 human diseases (Supplemental Table 3A). We found that DEGs in tau^0N3R^ brains overlapped significantly with modifiers associated with 37 disease classes (Figure 1E, Supplemental Table 3B). Strikingly, four of the top five most enriched were *Drosophila* models of neurodegenerative diseases—Huntington’s disease, tauopathy, amyotrophic lateral sclerosis and AD (Figure 1E, Supplemental Table 3B). This enrichment indicates that tau^0N3R^ triggers conserved neurodegenerative responses in fly brains. Consistent with our prior findings that tau^0N3R^-induced premature lethality in flies arose from elevated PLCβ–IP_3_R signaling (13), genes encoding several PLCβ-coupled GPCRs, both PLCβ paralogs, and IP_3_R were significantly upregulated in tau^0N3R^ brains (Figure 1F). Conversely, genes encoding proteins that terminate PLCβ–IP_3_R signaling were significantly downregulated (Figure 1F). Thus, functionally-relevant modifiers of neurodegeneration and tauopathy are embedded within the tau^0N3R^ transcriptomic signature.

Next, we asked whether human homologs of tau^0N3R^ DEGs are preferentially associated with neurological diseases using the DisGeNET repository, which catalogs gene–disease associations across 9,828 pathological conditions (18, 19). We extracted gene sets associated with 320 neurological, neurodegenerative, and neuromuscular diseases grouped into 32 broad categories (Supplemental Tables 4A-4B), and used the DIOPT ortholog finder (20) to identify 3,142 human genes with high-confidence homology to 2,757 DEGs (Supplemental Table 4C). Comparison of these DEG homologs with DisGeNET gene sets revealed a significant enrichment for genes associated with AD (Figure 1F, Supplemental Table 4D). Thus, AD-associated genes are disproportionately represented among the human homologs of tau^0N3R^-induced DEGs suggesting that observed transcriptomic changes might impinge upon pathways relevant to AD pathogenesis.

### Glycolysis emerges as a convergent pathway altered by both IP_3_R signaling and _tau0N3R_

Gene ontology and pathway analysis of the upregulated genes in *tau^0N3R^*-expressing brains revealed enrichment for neuronal excitability, neurotransmission and synaptic vesicle release, ion transport, and glycolysis (Supplemental Table 5A). Downregulated genes were enriched for proteasomal degradation, RNA processing, intracellular transport, cell adhesion, and neuronal differentiation (Supplemental Table 5B). These transcriptional changes are reminiscent of changes previously reported in various stages of human AD pathology (5).

Because the pathological effects of *tau^0N3R^* expression in *Drosophila* are mediated by increased *itpr* expression (13), we sought to identify the subset of the tau^0N3R^ transcriptomic signature that reflects the consequences of *itpr* overexpression. To this end, we performed bulk RNA-seq on larval brains isolated from animals ubiquitously overexpressing *itpr* (*tubulin>itpr*, Supplemental Tables 6A-6B). Differential expression analysis identified DEGs relative to each control (Figure S1A and Supplemental Tables 6C-6D). As expected, *itpr* itself was among the most significantly overexpressed genes in *tubulin>itpr* brains (Figure S1A). Intersecting DEG sets across the two controls pointed to 636 shared DEGs of which 627 (∼98.6%) displayed consistent directionality of gene expression changes (Figure S1B and Supplemental Table 6E). The resulting high confidence DEG set was comprised of 283 upregulated and 344 downregulated genes (Figure S1B and Supplemental Table 6E).

Comparison of *itpr* and *tau^0N3R^*-expressing transcriptomes revealed 479 shared DEGs, which constituted >75% of the DEGs from *itpr* overexpression (Figures S1C-S1D, Supplemental Table 6F). Not only were the DEGs highly concordant between the two genotypes, Fisher’s exact test indicated that the observed overlap was ∼15-fold higher than by chance. (Figures S1C-S1D). These data indicate that *itpr* overexpression DEGs represent a *bona fide* subset of the tau^0N3R^-induced transcriptomic response. Pathway analysis of the shared upregulated genes showed enrichment for glucose metabolism and glycolysis (Figure 1H, Supplemental Table 6G), whereas the shared downregulated genes were enriched for regulators of cell growth, differentiation, and morphogenesis (Figure S1E, Supplemental Table 6H). These findings implicate glycolysis as a convergent pathway altered by both IP_3_R and tau^0N3R^.

### Tau^0N3R^ drives coordinated upregulation of glycolysis components at both transcriptional and protein levels

Significantly upregulated genes in *tau^0N3R^*-expressing brains included those encoding transporters for glucose and trehalose (a glucose disaccharide, the preferred sugar source in insects (21, 22)), trehalase (converts trehalose to glucose (23)), all ten glycolytic enzymes, and lactate dehydrogenase (LDH) (Figure 2A). Conversely, genes encoding rate-limiting enzymes of oxidative branch of the pentose phosphate pathway (PPP), which directs glucose-6-phosphate away from glycolysis (24), were downregulated (Figure 2A). Likewise, genes encoding rate-limiting enzymes for gluconeogenesis, which drive flux in the direction opposite to that of glycolysis (25), were also repressed (Figure 2A). We observed no consistent changes in mitochondrial oxidative phosphorylation (OXPHOS) genes in tau^0N3R^ brains (Supplemental Tables 2C and 5A). These transcriptional changes suggest a metabolic commitment of glucose to glycolysis over competing pathways. To validate the RNA-seq findings, we quantified transcript levels in *tubulin>tau^0N3R^* brains relative to the *tubulin-GAL4*/+ and *UAS-tau^0N3R^*/+ control lines using quantitative RT-PCR. These analyses demonstrated consistent induction of glycolytic genes and *Ldh* in tau^0N3R^ brains relative to both parental controls (Figure 2B).

**Figure 2.**
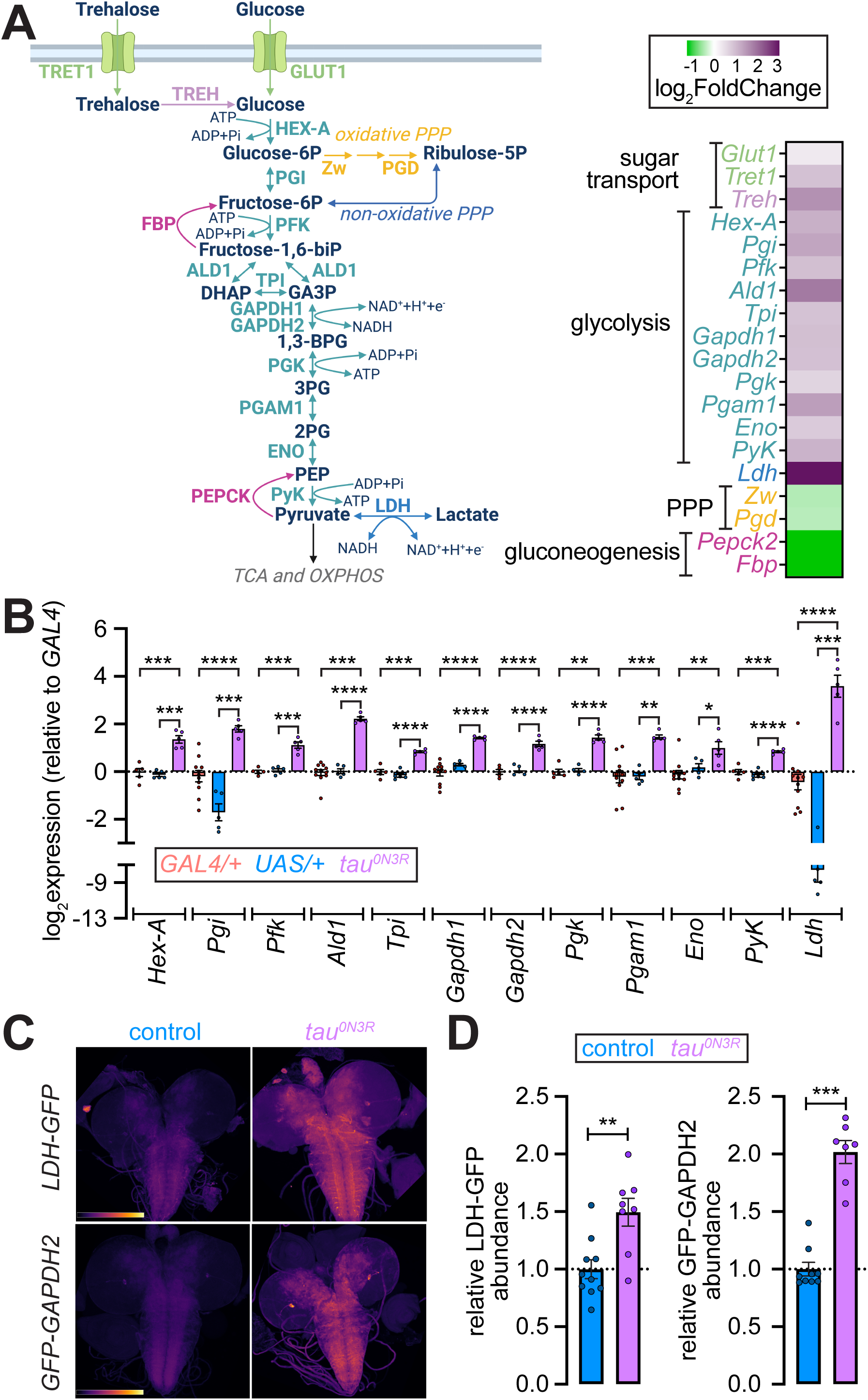
*Tau^0N3R^* expression drives coordinated upregulation of glycolytic genes and proteins in *Drosophila* brains and neurons. **(A)** Schematic of glucose and trehalose metabolism pathways (left), and heatmap (right) showing log_2_ fold changes of genes encoding sugar transporters, glycolytic enzymes, lactate dehydrogenase (*Ldh*), pentose phosphate pathway (PPP) enzymes, and gluconeogenic enzymes in *tau^0N3R^*-expressing brains. Color scale: magenta (upregulated), green (downregulated), gray (not significant). Genes encoding sugar transporters and trehalase (*Glut1*, *Tret1*, *Treh*), all glycolytic enzymes and LDH show upregulation. Genes encoding rate-limiting enzymes of the oxidative arm of the PPP (*Zw*, *Pgd*) and gluconeogenesis (*Pepck2*, *Fbp*) are downregulated. **(B)** Quantitative RT-PCR validation of glycolytic gene expression in larval brains. Log_2_ expression levels relative to *tubulin-GAL4*/+ control mean are shown. *Tubulin-GAL4*/+ (buff), *UAS-tau^0N3R^*/+ (blue), and *tubulin>tau^0N3R^* (magenta). Values represent mean ± SEM from n ≥ 3 biological replicates. Statistical comparisons were pairwise t-tests with Bonferroni post hoc corrections. \**P* < 0.05, \*\**P* < 0.01, \*\*\**P* < 0.001, \*\*\*\**P* < 0.0001. **(C)** Representative pseudocolored confocal images of larval brains from control (left column) and *tau^0N3R^*-expressing animals (*d42>tau^0N3R^*, right column) carrying endogenous GFP knock-in reporters for LDH (LDH-GFP, top row) or GAPDH2 (GFP-GAPDH2, bottom row). Pseudocolor represents intensity of GFP fluorescence. **(D)** Quantification of GFP fluorescence intensity from endogenous reporters in larval brains. Left, LDH-GFP; right, GFP-GAPDH2. All values are normalized to control means. Each point represents one brain; bars show mean ± SEM. Statistical comparison by unpaired t-test. \*\**P* < 0.01, \*\*\**P* < 0.001.

Next, we asked whether expression of *tau^0N3R^* in neurons alone increased glycolytic enzyme and LDH abundance. Because antibodies suitable for detecting *Drosophila* glycolytic enzymes are lacking, we turned to GFP-tagged lines in which *GFP* is fused to the endogenous coding sequences of *Ldh* and *Gapdh2*, and recapitulate native expression patterns (26–28). Expression of *tau^0N3R^* in glutamatergic neurons (using *d42-GAL4*) led to marked increases in abundance of LDH-GFP and GFP-GAPDH2 in brains (Figures 2C-2D), confirming that neuronal expression of *tau^0N3R^* results in the induction of glycolytic enzymes. Taken together, our data indicate that tau^0N3R^ led to coordinated upregulation of glycolysis components at both transcriptional and protein levels.

### Warburg-like metabolic reprogramming in tau^0N3R^ neurons

The coordinated induction of glycolytic genes in response to *tau^0N3R^* expression prompted us to probe the contributions of glycolysis and OXPHOS to neuronal ATP homeostasis. We coexpressed *tau^0N3R^* and the genetically-encoded ATP/ADP ratio sensor *PercevalHR* in glutamatergic neurons and conducted pH-corrected measurements in dissociated neurons, as described (13, 29, 30). In this assay, the overall decrease in ATP/ADP ratio upon the inhibition of OXPHOS or glycolysis reflects the contribution of that metabolic pathway to ATP production. Consistent with our prior work (29), inhibition of mitochondrial ATP synthase with oligomycin A (oligo A, 10 μM) induced a significant and sustained drop in ATP/ADP ratio in control neurons (*d42>PercevalHR*, Figures 3A and 3D). In tau^0N3R^ neurons, however, oligo A-evoked decline in ATP/ADP ratio was significantly attenuated, and appeared to partially recover toward baseline (Figures 3A and 3D). Conversely, 10 mM 2-deoxyglucose (2DG, a glycolysis inhibitor) lowered the ATP/ADP ratio to a significantly greater extent in tau^0N3R^ neurons than in controls (Figures 3B and 3D), revealing heighted dependence on glycolysis. Combined application of oligo A and 2DG reduced ATP/ADP ratios to the same extent in both genotypes (Figures 3C-3D). Notably, inclusion of 2DG eliminated the ratio recovery observed in oligo A-treated tau^0N3R^ neurons (Figures 3A and 3C), indicating that the partial resistance of the ATP/ADP ratio to oligo A in tau^0N3R^ neurons was the consequence of glycolytic flux. Together, these data point to comparable ATP demand and total ATP flux in both genotypes, but with a shift in the balance of ATP production toward glycolysis in tau^0N3R^—reminiscent of a Warburg-like metabolic reprogramming typical of cancer cells (31).

**Figure 3.**
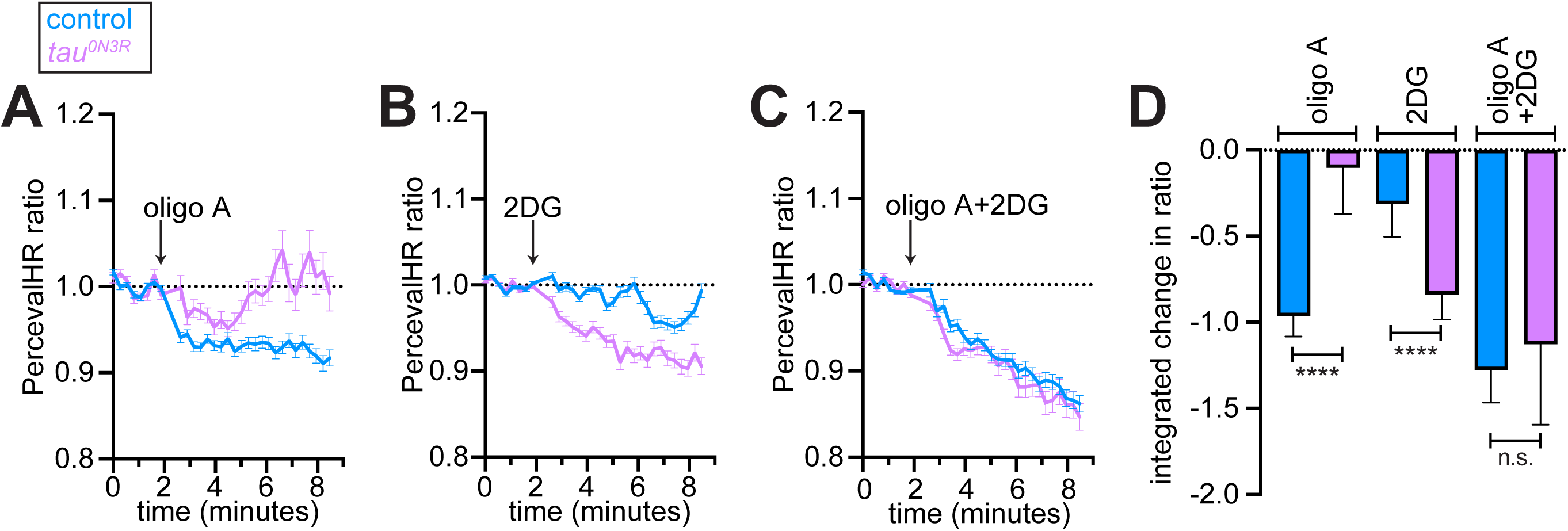
*Tau^0N3R^* expression shifts neuronal ATP production from oxidative phosphorylation to glycolysis. **(A–C)** Real-time ATP/ADP ratio measurements in dissociated glutamatergic neurons coexpressing *tau^0N3R^* and the genetically encoded ATP/ADP sensor *PercevalHR*. Traces show pH-corrected PercevalHR ratio normalized to baseline (dotted line at 1.0) for control neurons (*d42>PercevalHR*, blue) and *tau^0N3R^*-expressing neurons (*d42>PercevalHR+tau^0N3R^*, magenta). **(A)** Response to 10 μM oligomycin A (oligo A). **(B)** Response to 10 mM 2-deoxyglucose (2DG). **(C)** Response to the combined application of 10 μM oligomycin A and 10 mM 2DG. Data are mean ± SEM from n > 60 neurons per genotype across ≥ 3 independent experiments. **(D)** Quantification of integrated ATP/ADP ratio changes following pathway inhibition. Bars represent median ± 95% confidence intervals. \*\*\*\**P* < 0.0001, n.s., not significant, Mann-Whitney tests.

To corroborate the metabolic shift inferred above, we performed Seahorse metabolic flux analyses on brains expressing *tau^0N3R^* in glutamatergic neurons (*d42>tau^0N3R^*) and the respective controls (*d42-GAL4*/+). First, we used the glycolytic rate assay (Supplemental Table 7A) to determine the glycolytic response to acute inhibition of OXPHOS with rotenone and antimycin A. Under standard assay conditions (glucose, pyruvate and glutamine in assay media), both genotypes achieved comparable maximal extracellular acidification rate (ECAR) in response to OXPHOS inhibition (Figure 4A, left and Supplemental Table 7B), indicating that the upper limit of glycolytic capacity was not altered in tau^0N3R^ brains. However, the glycolytic reserve—the difference between maximal and basal ECAR—was significantly reduced in tau^0N3R^ brains (Figure 4A, left and 4B, Supplemental Table 7B). These data indicate that while the glycolytic ceiling remained unchanged in tau^0N3R^ brains these brains operated significantly closer to that ceiling at rest, consistent with a shift toward glycolytic ATP sourcing.

**Figure 4.**
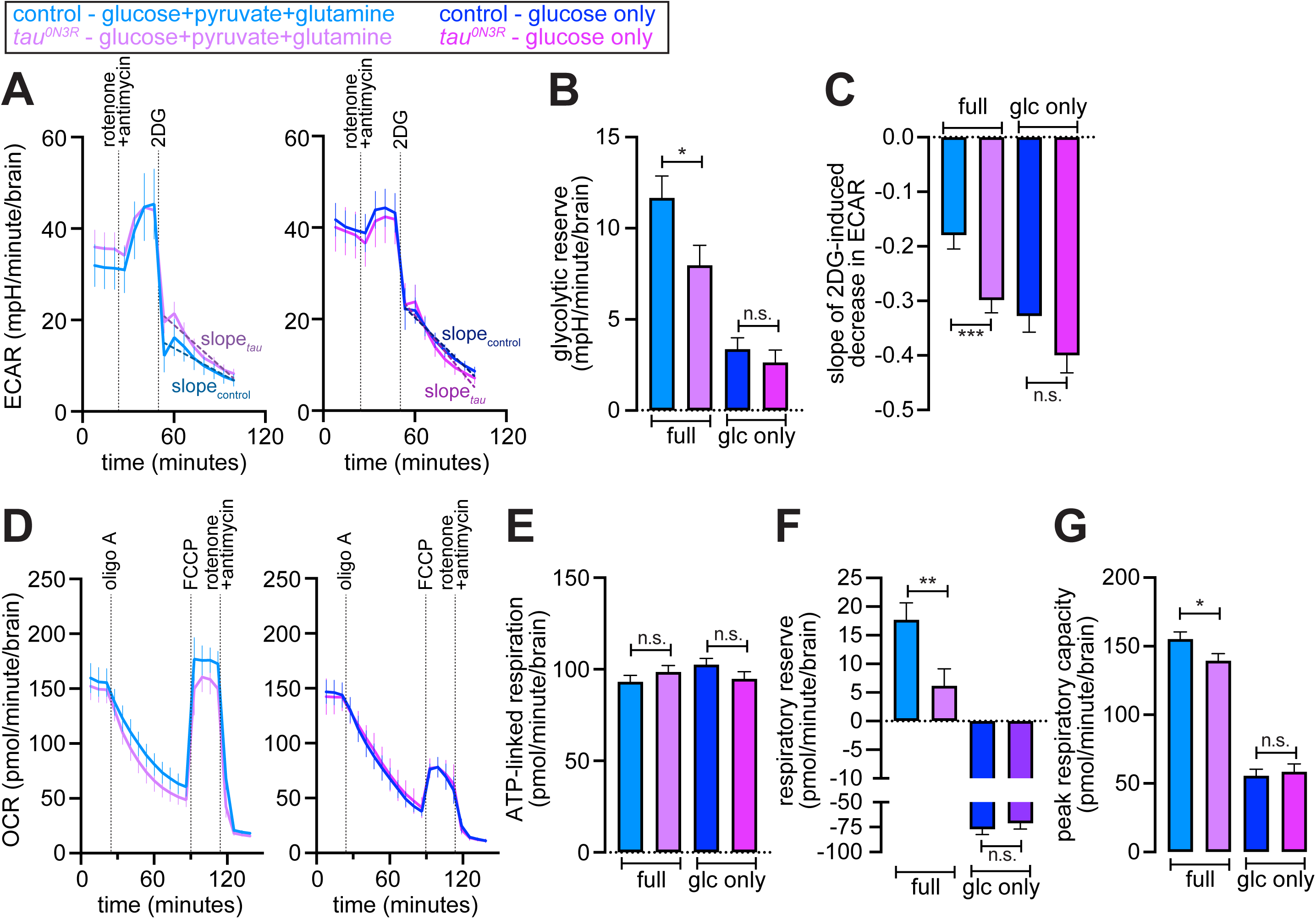
Expression of *tau^0N3R^* in glutamatergic neurons results in elevated basal glycolysis with reduced glycolytic and respiratory reserves. **(A)** Seahorse extracellular acidification rate (ECAR) measurements on *ex vivo* larval brains expressing *tau^0N3R^* in glutamatergic neurons (*d42>tau^0N3R^*, magenta) and controls (*d42-GAL4*/+, blue). Left, full medium containing 10 mM glucose, 1 mM pyruvate, and 2 mM glutamine. Right, 10 mM glucose-only medium (pyruvate and glutamine omitted). Sequential additions of rotenone + antimycin (10 μM each) and 2DG (200 mM) are indicated. Data are mean ± SEM from n = 5 control and n = 6 tau^0N3R^ brains (full medium); n = 6 control and n = 5 tau^0N3R^ brains (glucose-only). **(B)** Quantification of glycolytic reserve calculated as the increase in ECAR following rotenone + antimycin addition in the indicated media. Values represent mean ± standard error, \**P* < 0.05, n.s., not significant, linear mixed-effects model (LMM) accounting for variation between individual wells and experimental batches (ECAR ∼ condition * group + (1 | well) + (1 | batch)). **(C)** Rate of 2DG-induced ECAR suppression calculated as the slope of ECAR decline after 2DG addition in the indicated media. Values represent mean ± standard error, \*\*\**P* < 0.001, n.s., not significant, LMM model accounting for variation between individual wells and experimental batches (ECAR ∼ time * group + (1 | well) + (1 | batch)). **(D)** Seahorse oxygen consumption rate (OCR) traces from mitochondrial stress test showing sequential additions of oligomycin A (oligo A, 10 μM), FCCP (3 μM), and rotenone + antimycin (10 μM each). Left, full medium. Right, glucose-only medium. Data are mean ± SEM from n = 7 control and n = 7 tau^0N3R^ brains (full medium) or n = 4 control and n = 3 tau^0N3R^ brains (glucose-only). **(E)** Quantification of ATP-linked respiration, calculated as the reduction in OCR following oligomycin A addition in the indicated media. Values represent mean ± standard error, n.s., not significant, LMM model accounting for variation between individual wells and experimental batches (OCR ∼ condition * group + (1 | well) + (1 | batch)). **(E)** Quantification of respiratory reserve calculated as the increase in OCR from baseline following FCCP addition in the indicated media. Values represent mean ± standard error, n.s., not significant, LMM model accounting for variation between individual wells and experimental batches (OCR ∼ condition * group + (1 | well) + (1 | batch)). **(G)** Quantification of peak respiratory capacity measured as maximal OCR following FCCP addition in the indicated media. Values represent mean ± standard error, \**P* < 0.05, n.s., not significant, LMM model accounting for variation between individual wells and experimental batches (OCR ∼ condition * group + (1 | well) + (1 | batch)).

Omission of pyruvate and glutamine from assay media (glucose-only condition), which limits flux through the tricarboxylic acid (TCA) cycle and results in the homeostatic augmentation of glycolysis, similarly had no impact on maximal ECAR in control and tau^0N3R^ brains (Figure 4A, right and Supplemental Table 7B). As expected, glucose-only condition increased basal glycolysis and reduced glycolytic reserves in both genotypes (Figures 4A, right and 4B, Supplemental Table 7B). Notably, under the substrate-limited conditions, the significant difference in glycolytic reserve between the genotypes observed in full-nutrient medium was abolished (Figures 4A-4B, Supplemental Table 7B). This convergence indicates that tau^0N3R^ brains constitutively operate closer to their glycolytic ceiling, and that this difference is masked when control brains are forced into a similar high-glycolysis state by removal of TCA substrates.

Application of 2DG led to rapid decreases in ECAR across genotypes and media conditions (Figure 4A). The slope of a linear fit to ECAR values post 2DG application (dashed lines superimposed on traces, Figure 4A) was significantly higher in tau^0N3R^ brains relative controls under full-nutrient conditions (Figures 4A, left and 4C, Supplemental Table 7B). In glucose-only medium, 2DG-induced slopes increased in both genotypes (Figure 4A, right and 4C, Supplemental Table 7B). As with glycolytic reserve, the difference in slopes between genotypes was abolished under glucose-only conditions (Figures 4A and 4C, Supplemental Tables 7A-7B). Together, these data indicate heightened engagement of glycolysis in tau^0N3R^ that is obscured when the control brains are forced closer to their maximal glycolysis.

Next, we performed the mitochondrial stress test to assess oxygen consumption rates (OCR) in tau^0N3R^ and control brains (Supplemental Table 7C). Both basal OCR and ATP-linked OCR (oligo A-sensitive OCR) were unchanged between genotypes under either media condition (Figures 4D-4E, Supplemental Table 7D). Therefore, mitochondrial ATP production under resting energetic conditions is preserved in tau^0N3R^ brains. In contrast, FCCP-induced maximal OCR (i.e., maximal OXPHOS capacity) was lower in tau^0N3R^ brains under full nutrient conditions (Figure 4D, left). Accordingly, both respiratory reserve (difference between maximal and basal OCR) and peak respiratory capacity (difference between maximal OCR and OCR after rotenone and antimycin A treatment) were significantly lower in tau^0N3R^ (Figures 4F-4G, Supplemental Table 7D). Restricting the supply of TCA-derived reducing equivalents by omission of pyruvate and glutamine from the assay medium lowered maximal OCR, respiratory reserve, and peak respiratory capacity in both genotypes (Figures 4D, right and 4F-4G). Under these substrate-restricted conditions, the genotype differences observed in full-nutrient medium were abolished (Figures 4F-4G), mirroring the glycolytic reserve measurements.

Collectively, these results indicate that tau^0N3R^ brains exhibit elevated basal glycolysis and reduced glycolytic reserve, accompanied by a reduction in maximal but not basal mitochondrial respiration. The preservation of basal and ATP-linked OCR, together with the reduced FCCP-stimulated maximal OCR in tau^0N3R^ is consistent with a limitation in the ability to increase oxidative flux rather than a defect in the core OXPHOS machinery. The loss of genotype differences under glucose-only conditions indicates that tau^0N3R^ brains constitutively operate in a metabolic state resembling TCA substrate limitation, which can be masked by forcing controls into a similar substrate-restricted state. Together, these data demonstrate that *tau^0N3R^* expression triggers a Warburg-like shift prioritizing glycolysis for maintaining ATP homeostasis despite preserved basal oxidative metabolism.

### Tau^0N3R^-induced glucose hypermetabolism and elevated lactate production lead to accelerated biological aging

Having established that tau^0N3R^ drives Warburg-like shift to glycolytic metabolism, we next investigated whether the increases in glycolytic gene expression contributes to the premature lethality caused by *tau^0N3R^* expression in adult glutamatergic neurons (Figure S2A, Supplemental Table 8A) (13). We first assessed whether glycolytic genes are required for viability by testing RNAi lines targeting enzymes across the glycolytic pathway (Figure 5A). Knockdown of genes encoding aldolase (ALD1), enolase (ENO) or the rate-limiting enzymes of glycolysis—phosphofructokinase (PFK) and pyruvate kinase (PyK)—led to developmental lethality, precluding phenotypic analysis in adults (lethal genes marked in red, Figure 5A). In contrast, knockdown of trehalose transporter (*Tret1*), hexokinase (*Hex-A*), *Gapdh1*, phosphoglycerate kinase (*Pgk*), and phosphoglycerate mutase (*Pgam1*) had no effects on development or animal viability. Remarkably, knockdown of each of these genes in *tau^0N3R^*-expressing glutamatergic neurons suppressed tau^0N3R^-induced premature lethality, restoring adult lifespan towards control levels (Figures 5B-5H, Supplemental Table 8A). For *Gapdh1* and *Pgk*, independent RNAi lines yielded consistent rescue, ruling out off-target effects (Figures 5D-5G, Supplemental Table 8A). Knockdown of *Ldh* also restored the lifespan of *tau^0N3R^*-expressing animals (Figure 5I, Supplemental Table 8A).

**Figure 5.**
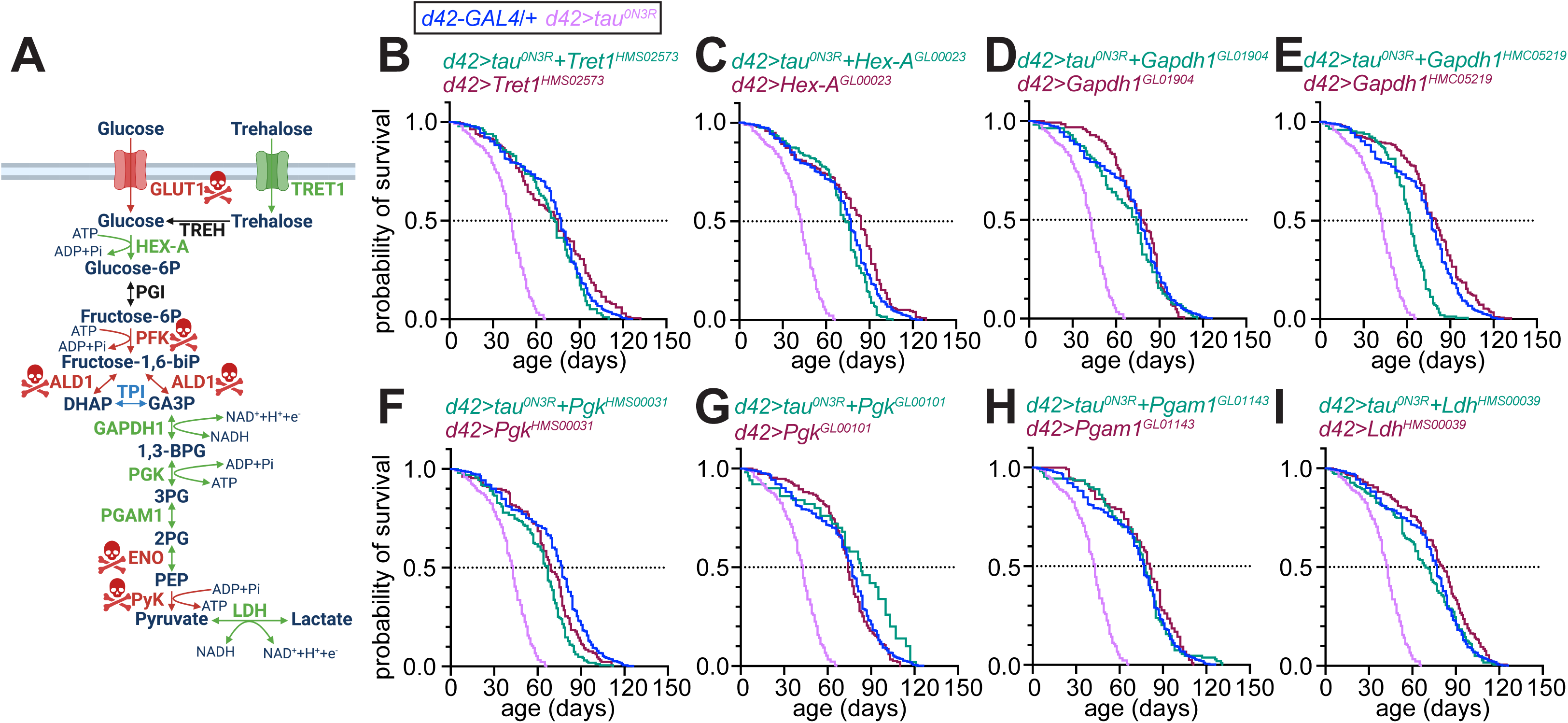
RNAi-mediated knockdown of glycolytic genes and lactate dehydrogenase rescues tau^0N3R^-induced premature lethality. **(A)** Schematic of glucose and trehalose metabolism. Knockdown of genes marked with skull-and-crossbones symbols in red cause developmental lethality. Knockdown of genes shown in green permit viability, and rescued the effects of tau^0N3R^ on adult lifespan. Knockdown of the gene shown in blue permits viability, but did not rescue the effects of tau^0N3R^ on adult lifespan. Genes marked in black were not examined. **(B–I)** Kaplan-Meier survival curves for adult flies of the indicated genotypes. Dotted line indicates 50% survival. See Supplemental Table 8A for sample sizes of all genotypes.

Using qRT-PCR, we confirmed that each RNAi line that restored animal lifespan also significantly reduced the mRNA levels of its target gene (Figure S2B). An exception was *Tpi*, whose knockdown produced no lifespan extension despite significant reduction in transcript levels (Figures S2B-S2C, Supplemental Table 8A), suggesting limited contribution of this enzyme in sustaining glycolytic flux in *tau^0N3R^*-expressing glutamatergic neurons. Importantly, absence of a suppressive effect associated with the coexpression of *Tpi* RNAi also argues that the rescue of lifespan by the other glycolytic RNAi lines were not the consequences of GAL4 dilution stemming from the presence of two *UAS* lines. In case of *Gapdh1*, the RNAi line achieving stronger transcript knockdown produced a correspondingly greater rescue of lifespan (Figures 5D-5E and S2B, Supplemental Table 8A), pointing to a dose dependent relationship between glycolytic gene expression and toxicity. Collectively, these findings indicate that elevated glycolysis and lactate production are key mediators of tau^0N3R^-induced premature lethality in *Drosophila*.

To determine whether shorter lifespan in flies expressing *tau^0N3R^* in glutamatergic neurons reflected increased baseline mortality or accelerated aging rates, we applied Gompertz mortality models to the fly survival data. The Gompertz model parameterizes mortality hazard as h(t) = α·exp(βt), where α represents baseline mortality and β represents the aging rate—the exponential increase in mortality risk with age (15). We first validated the Gompertz assumption by examining empirical hazard estimates (Supplemental Table 8B). Log-hazard showed strong linear relationships with time (R^2^∼0.9 across genotypes, Figure 6A and Supplemental Table 8C), confirming the appropriateness of parametric Gompertz modeling.

**Figure 6.**
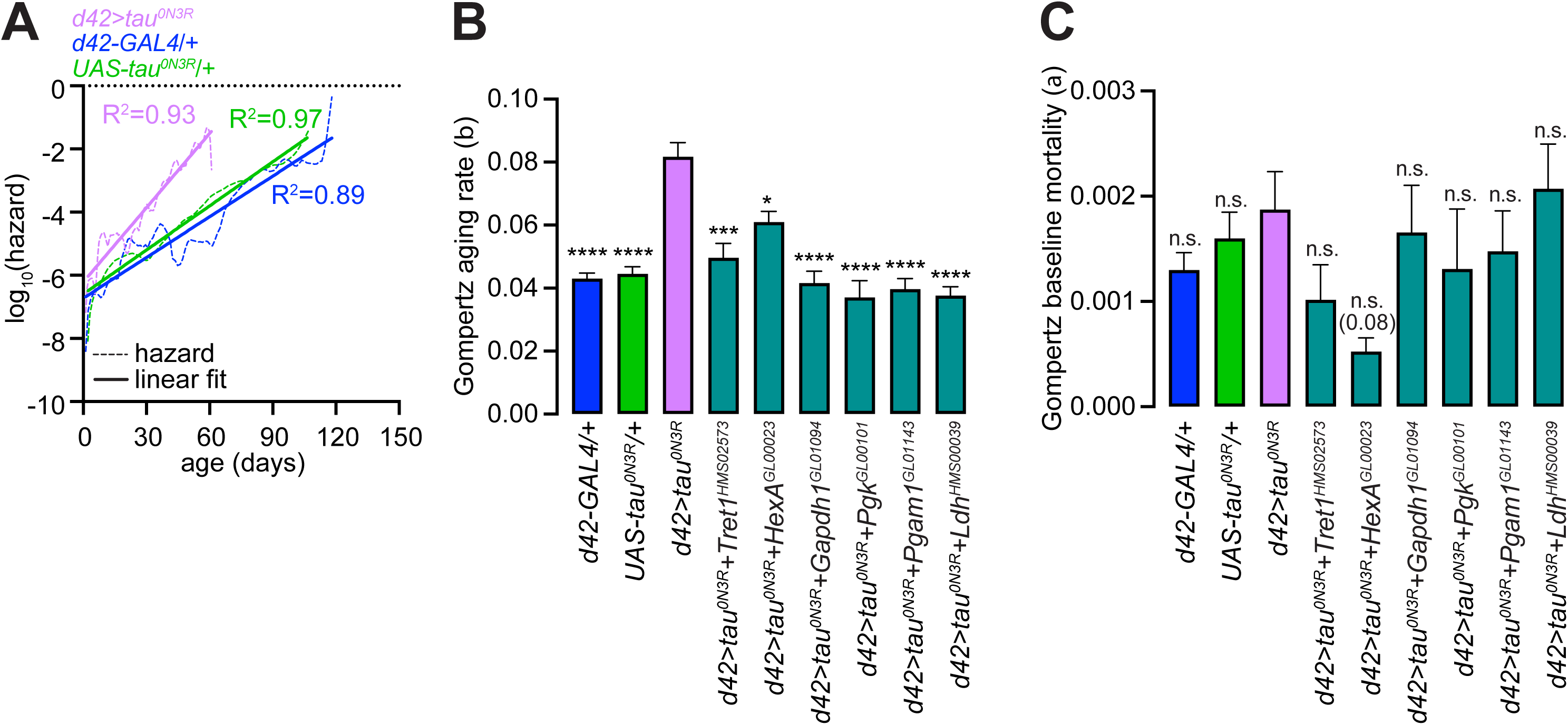
Tau^0N3R^ specifically increases aging rate, which is normalized by glycolytic gene knockdown. **(A)** Validation of Gompertz mortality model assumption in the indicated genotypes. Empirical hazard estimates (dashed lines), linear fits (solid lines), and R^2^ values for fits are shown. **(B-C)** Gompertz aging rate parameter **(B)** and baseline mortalities **(C)** in the indicated genotypes. Values represent mean ± standard error from Gompertz model fits on lifespan data. All statistical comparisons were made with *d42>tau^0N3R^* flies. \*\*\*\**P* < 0.0001, \*\*\**P* < 0.001, \**P* < 0.05, n.s., not significant, Wald z-test on pairwise parameter differences with Holm post hoc correction for multiple comparisons.

Upon fitting the survival data to the Gompertz model, we found that relative to the *UAS* and *GAL4* controls, expression of *tau^0N3R^* in glutamatergic neurons significantly increased the aging rate parameter (β) without significantly affecting baseline mortality (α) (Figures 6B-6C and Supplemental Table 8C). Therefore, tau^0N3R^ accelerates the aging process itself rather than imposing a fixed mortality cost. Knockdown of key glycolytic genes or *Ldh* in *tau^0N3R^*-expressing neurons normalized the aging rate parameter without affecting baseline mortality (Figures 6B-6C and Supplemental Table 8C). Knockdown of these glycolytic genes in glutamatergic neurons of control animals did not lower aging rate nor did it induce consistent changes in baseline mortality (Figures S3A-S3B and Supplemental Table 8C). Together, these data demonstrate that glutamatergic expression of human *tau^0N3R^* specifically increases the aging rate via the augmentation of glucose metabolism and lactate production.

## DISCUSSION

We show that human *tau^0N3R^* expression in *Drosophila* neurons triggers Warburg-like metabolic shift towards increased glycolysis, with decreased reliance on OXPHOS for ATP production. This metabolic reprogramming is driven by both transcript and protein level changes in glycolytic enzymes and LDH. Similar increases in glycolytic components and LDH have been reported in independent studies of single-cell transcriptomic changes in cortical neurons from patients with early-onset AD (7), in proteomic studies conducted on AD patients with high burden of tau pathology (5), in vulnerable brain regions of the PS19 mouse model of tauopathy/AD (4), and in cultured cells expressing human *tau* (32). Most compellingly, direct conversion of sporadic AD patient fibroblasts into induced neurons that maintain epigenetic features of donor age recapitulated the Warburg-like metabolic switch to aerobic glycolysis, accompanied by significant increase in the expression of the gene encoding lactate dehydrogenase (*LDHA*) (8, 9). The induction of glycolytic metabolism in induced AD neurons was mediated by STAT3 and HIF1α signaling, and promoted neuronal vulnerability (8, 12). The convergence of our findings in *Drosophila* with multiple orthogonal datasets spanning transcriptomics, proteomics, mouse models and patient-derived neurons validates the relevance of our model to human AD pathogenesis, and establishes that *tau^0N3R^* expression in fly neurons is sufficient to drive metabolic reprogramming reminiscent of AD neuropathology in humans.

Critically, our study extends beyond correlation to reveal the functional consequences of neuronal metabolic reprogramming. We demonstrate that the increase in neuronal glycolysis in tau^0N3R^ neurons exacts a cost in the form of accelerated biological aging. Genetic suppression of glycolysis or Ldh specifically in *tau^0N3R^*-expressing neurons restored normal aging kinetics, proving that metabolic reprogramming is neither a passive reflection of the consequences of neuronal damage nor does it merely correlate with toxicity. Instead, the Warburg-like metabolic shift mechanistically drives tau^0N3R^-induced neurotoxicity. This causal demonstration— unattainable in human correlative studies—establishes that the altered metabolic strategy in *tau^0N3R^*-expressing neurons, rather than ATP levels *per se*, determines aging rate. This idea is consistent with the notion of antagonistic pleiotropy during aging (33), whereby an adaptive response to acute stress (maintenance of ATP) produces detrimental long-term consequences (faster aging).

### Metabolic dysfunction in a model of tauopathy

#### What might drive glucose hypermetabolism in tau^0N3R^ neurons?

Seahorse metabolic flux analyses revealed that tau^0N3R^ does not impair basal OXPHOS but selectively reduces maximal respiratory capacity. This pattern is consistent with a limitation in the ability to *upregulate* mitochondrial oxidative flux rather than a deficit in the OXPHOS machinery itself. The abolition of genotype differences under glucose-only conditions further supports the interpretation that tau^0N3R^ neurons operate in a metabolic state that resembles substrate limitation at the level of TCA cycle input, such that forcing controls into a similar substrate-restricted state masks the phenotype.

One plausible upstream contributor is altered mitochondrial Ca^2+^ handling. We previously showed that neurons expressing *tau^0N3R^* exhibit diminished mitochondrial Ca^2+^ uptake (13). Because matrix free Ca^2+^ activated key TCA dehydrogenases (34), diminished mitochondrial Ca^2+^ uptake would be expected to constrain the ability of mitochondria to increase TCA cycle flux during elevated energetic demand, thereby lowering maximal respiratory capacity despite preserved basal respiration. In this framework, the glycolytic shift observed here is not secondary to mitochondrial failure, but could arise from a reduced capacity to upregulate oxidative metabolism when needed.

### Tauopathies as diseases of accelerated aging

Our application of Gompertz mortality modeling to dissect mechanisms underlying shorter lifespan in a fly model of tauopathy provides a quantitative framework of broad utility in neurodegeneration models with early mortality. Conventional metrics such as median and maximum lifespans conflate the effects of baseline mortality and aging rates. By decomposing survival data using Gompertz modeling, we demonstrate that tau^0N3R^ specifically accelerates aging rates (Gompertz β parameter) without increasing baseline mortality (Gompertz α parameter). These results agree with reports of accelerated aging trajectories in brains of AD patients as determined using DNA methylation-based aging clocks (35–37). Importantly, we show that knockdown of glycolytic genes or *Ldh* in glutamatergic neurons expressing *tau^0N3R^* specifically restored aging rate without affecting baseline mortality. This reversal of the Gompertz β parameter implies a causal role for glycolysis in tau^0N3R^-induced shortening of lifespan and not a general health benefit unrelated to the effects tau^0N3R^ on aging. Our findings agree with prior reports of a shift to Warburg-like aerobic glycolysis in hippocampal neurons during normal aging in mice, and the contribution of this metabolic change to age-related cognitive impairment and synapse loss (38).

#### Why would glycolytic upregulation accelerate aging?

Several non-mutually exclusive mechanisms are plausible. First, glycolysis- and LDH-dependent lactate production may create stress through extracellular acidification or direct lactate-mediated signaling. Lactate is a potent signaling molecule that contributes to progressive toxicity in a mouse model of AD via its impacts on histone modification and gene expression (39–41). More pertinently, overexpression of *Ldh* in *Drosophila* neurons promotes age-dependent neurodegeneration and shortens animal lifespan, whereas *Ldh* reduction delays neurodegeneration (42)—findings that are highly concordant with ours. Second, depletion of NAD^+^ pools due to excessive glycolysis could impair NAD^+^-dependent longevity pathways involving sirtuins and DNA repair enzymes (43, 44). Indeed, by relying on glia-derived metabolites that fuel the TCA cycle, healthy mature neurons keep their reliance on glycolysis low in order to preserve cytosolic NAD^+^ levels (43). Third, elevated glycolysis could compromise flux through the PPP, the primary source of NADPH for antioxidant defense (24). Transcriptional downregulation of genes encoding rate-limiting PPP enzymes would contribute to lower flux through the pathway. In turn, reduced NADPH production would impair regeneration of reduced glutathione and thioredoxin, thereby diminishing neurons’ capacity to neutralize reactive oxygen species—an established driver of aging. Given neurons’ particular vulnerability to oxidative stress and redox imbalance, these cells typically exhibit preference for PPP over glycolysis (43, 45).

### Limitations and future directions

Our study has a few limitations of note. We characterized metabolic changes in larval brains and neurons, while the aging phenotype manifested in adults. Although the restoration of aging rate by knockdown of glycolytic genes identified in larval neurons suggests that early-life metabolic remodeling predicts adult outcomes, future work should directly measure glycolytic engagement in adult neurons *in vivo*.

Second, our experiments were carried out in a single model organism and with one human tau isoform. Significant overlap between our transcriptional signature and human AD-associated genes suggests conservation, but direct cross-species comparisons will be necessary. Recent studies demonstrating that the expression of the 0N4R isoform of human *tau* in *Drosophila* neurons accelerates rates of aging in peripheral tissues as determined by snRNA-seq-derived aging clocks (46), demonstrates that neurons expressing a different *tau* isoform produce systemic signals that promote organism-wide aging. Whether the glycolytic phenotype we describe contributes to such systemic influence is an important question for future study.

Finally, our analyses focused on neurons, but glial cells are increasingly recognized as key regulators of brain metabolism in both normal physiology and AD. Glia are major sources of lactate and metabolic support to neurons (47, 48). Whether tau-induced metabolic reprogramming is reinforced, buffered, or even initiated by changes in glial metabolism or glia–neuron metabolic coupling remains unknown. Future studies incorporating cell type–specific metabolic profiling will be required to determine how neuron–glia signaling shapes the metabolic phenotype observed here.

## ACKNOWLEDGEMENTS

We thank the Bloomington *Drosophila* Stock Center for fly stocks. Live-cell fluorescence microscopy and image analysis was performed at the Center for Advanced Microscopy, a Nikon Center of Excellence, in the Department of Integrative Biology & Pharmacology at McGovern Medical School, UTHealth Houston. We are grateful to Dr. Neal Waxham for reading the manuscript and providing feedback. This work was supported by National Institutes of Health (NIH) grants R01AG069076, R01AG072176, and R21AG087381 to K.V.

## AUTHOR CONTRIBUTIONS

R.G. and K.V. designed the research; R.G., E.R., H.M. and M.S.P. performed the experiments; R.G. and K.V. analyzed data and wrote the manuscript.

## DECLARATION OF INTERESTS

The authors declare no competing interests.

## MATERIALS AND METHODS

### *Drosophila* husbandry and fly stocks

Flies were reared at room temperature on standard fly food (per 1 L: 95 g agar, 275 g brewer’s yeast, 520 g cornmeal, 110 g sugar, 45 g propionic acid, and 36 g Tegosept). Stocks obtained from the Bloomington *Drosophila* Stock Center included *w^1118^*, *d42-GAL4* (49), *tubulin-GAL4* (50), *hs-GAL4* (51), *UAS-itpr* (52), *UAS-tau^0N3R^* (stock number 181) (17), *LDH-GFP* (26), *GFP-GAPDH2* (28), and all RNAi lines (*UAS-Glut1^HMS02152^*, *UAS-Tret1^HMS02573^*, *UAS-Hex-A^GL00023^*, *UAS-Pfk^HMS01324^*, *UAS-Ald1^HMC06146^*, *UAS-Eno^HMC05514^*, *UAS-Gapdh1^GL01904^*, *UAS-Gapdh1^HMC05219^*, *UAS-Pgk^HMS00031^*, *UAS-Pgk^GL00101^*, *UAS-Pgam1^GL01143^*, *UAS-Pyk^HMC04605^*, *UAS-Ldh^HMS00039^*). *UAS-PercevalHR* was described previously (13).

### Whole RNA sequencing and data analyses

#### RNA purification and sequencing

3^rd^ instar larval brains were dissected in ice-cold RIPA buffer. Total RNA was extracted using the RNeasy Mini Kit (Qiagen) following the manufacturer’s protocol and eluted in RNase-free water. Each biological replicate was comprised of RNA extracted from 60-70 brains pooled together. We prepared six biological replicates per genotype. RNA concentration and purity were quantified using a NanoDrop spectrophotometer (Thermo Scientific). High-quality RNA samples were submitted to Novogene for library preparation and whole transcriptome RNA sequencing. Paired-end sequencing was performed on an Illumina NovaSeq PE150 platform with a target depth of 20-30 million reads per sample.

#### Read alignment and quantification

Raw sequencing reads were processed using the Rsubread package (53) in R. A reference index was built from the *Drosophila melanogaster* reference genome (BDGP6.32). Paired-end reads from each sample were aligned to the reference genome. Gene-level read raw counts were generated with a *Drosophila melanogaster*-specific gene annotation file (BDGP6.32.107 from Ensembl). Paired-end information was preserved during counting. For normalization of gene expression, we calculated transcripts per million (TPM) values to account for both sequencing depth and gene length biases. Gene lengths were extracted from an annotation file, and were calculated as the sum of exon lengths for each gene. Raw counts were first normalized by gene length (reads per kilobase, RPK), then scaled to the total library size to generate TPM values using the formula: TPM = (RPK / sum of all RPK values) × 10^6^.

#### Principal component analysis (PCA) and sample clustering

PCA was performed on TPM-normalized expression values. Genes with zero expression across all samples were removed prior to analysis. Expression values were centered and scaled using z-score transformation. K-means clustering (k = 2) was applied. Clustering results were visualized with sample labels and confidence ellipses using the ggfortify package (54) in R.

#### Differential expression analysis

Differential gene expression analysis was performed using DESeq2 package (55) in R. Raw count matrices were filtered to only retain genes with total counts > 1 across all samples. Samples were grouped into control and treatment conditions. The DESeq2 model was fit using default parameters. Differential expression between treatment and control groups was assessed using the Wald test, with Cook’s distance outlier detection and independent filtering disabled to retain all genes for downstream analyses. Gene-level fold changes, *P*-values, and adjusted *P*-values were calculated using the Benjamini-Hochberg procedure for multiple testing correction. Differentially expressed genes were identified using an adjusted *P*-value threshold of 0.05.

### Quantitative RT-PCR

#### cDNA synthesis and qRT-PCR

1 μg total RNA was reverse transcribed into cDNA using the High-Capacity RT kit (Thermo Fisher). Quantitative RT-PCR was performed using PowerUp SYBR Green PCR Master Mix (Thermo Fisher) on a CFX384 thermocycler (Bio-Rad). Reactions were performed in triplicates. The mRNA abundance of each gene was normalized to the housekeeping gene *rp49* using the 2^-ΔΔCt^ method. Supplemental Table 9 provides primer sequences.

#### RNA extraction from larval brains

For analyses of gene expression in brain, total RNA was extracted from the 3^rd^ instar larval brains dissected in ice-cold PBS and homogenized in RIPA buffer using the RNeasy Mini Kit (Qiagen) according to the manufacturer’s protocol. RNA concentration and purity were assessed using a NanoDrop spectrophotometer (Thermo Scientific).

#### RNAi validation

RNAi-mediated gene knockdown efficiency was quantified as described (13, 29). Briefly, flies expressing RNAi transgenes under the control of a heat shock-inducible promoter (*hs-GAL4*) were subjected to heat shock treatment (1 hour at 37°C) on two consecutive days. Total RNA was extracted 24 hours after the second heat shock using the RNeasy mini kit (Qiagen) following the manufacturer’s instructions. Gene knockdown efficiency was calculated relative to *hs-GAL4/+* controls subjected to the same heat shock.

### Brain immunostaining and image analysis

#### Tissue preparation and immunofluorescence

3^rd^ instar larval brains were dissected in ice-cold PBS and fixed in 4% paraformaldehyde for 1 hour at room temperature. Fixed tissues were washed three times for 5 minutes each in PBS containing 0.1% Triton X-100 (PBS-T). Brains were blocked and permeabilized in 5% normal donkey serum (Sigma-Aldrich) in PBS-T, then incubated overnight at 4°C with primary antibodies— mouse monoclonal anti-tau (clone T46, 1:1000; Invitrogen) and rabbit monoclonal anti-GFP (1:1000; Cell Signaling Technology). Following three washes in PBS-T, samples were incubated with secondary antibodies for 1 hour at room temperature—Alexa Fluor 568 goat anti-mouse and Alexa Fluor 488 goat anti-rabbit (both 1:500; Invitrogen). After three final washes in PBS-T, brains were mounted on glass slides using Fluoromount-G with DAPI (Southern Biotech).

#### Confocal microscopy and image quantification

Z-stack images of immunostained brains were acquired using a Nikon A1R confocal microscope with a 20X objective. Imaging parameters such as laser power, gain, and pinhole settings were maintained constant across all samples. Image analysis was performed using Fiji/ImageJ software. For quantification of GFP signal intensity, maximum intensity projections were generated from z-stacks. A region of interest (ROI) was manually drawn around each brain to exclude non-brain tissue. Background fluorescence was subtracted using the rolling ball method (radius = 50 pixels), and integrated GFP fluorescence intensity was measured for each brain. Statistical comparisons were performed using GraphPad Prism.

### Dissociation and culture of *Drosophila* neurons

We dissociated primary neurons from 3^rd^ instar brains as described previously (13, 29, 30). Briefly, larvae were surface-sterilized by brief immersion in 70% ethanol followed by washing in sterile distilled water. Brains were dissected in filtered Schneider’s medium (Sigma-Aldrich) supplemented with 10% heat-inactivated fetal bovine serum (FBS), 1% antibiotic-antimycotic solution (Sigma-Aldrich), and 50 μg/mL bovine insulin (Sigma-Aldrich). For enzymatic dissociation, brains were transferred to sterile-filtered HL-3 (70 mM NaCl, 5 mM KCl, 1 mM CaCl_2_, 20 mM MgCl_2_, 10 mM NaHCO_3_, 115 mM sucrose, 5 mM trehalose, 5 mM HEPES; pH 7.2) containing 0.423 mM L-cysteine (MilliporeSigma) and 5 U/mL papain (Sigma-Aldrich). Enzymatic digestion proceeded for 30 minutes at 25°C. Following papain treatment, brains were washed once with complete Schneider’s medium and transferred to 1.5 mL microcentrifuge tubes for mechanical dissociation by gentle trituration. Dissociated neurons were plated on 35 mm glass-bottom dishes (Cellvis) pre-coated with 0.5 mg/mL concanavalin A (Sigma-Aldrich). Neuronal cultures were maintained at 25°C in a humidified chamber for 4 days, with daily medium replacement to prevent yeast contamination.

### Live imaging of *Drosophila* neurons and image analysis

#### Fluorescence imaging

Dissociated *Drosophila* neurons were imaged on day 5 post-plating as described (30). Briefly, neurons were incubated in Schneider’s media containing pHrodo-Red AM (ThermoFisher) for 30 minutes at 25°C. Following two washes with fresh Schneider’s medium, neurons were equilibrated in HL-3 for 10 minutes before imaging. Live-cell imaging of neurons was performed using a Nikon A1R laser-scanning confocal microscope equipped with a 40× Plan Fluor oil immersion objective and the Perfect Focus System for focal drift correction. PercevalHR fluorescence was acquired using dual excitation at 405 nm and 488 nm with emission collected at 535 nm. The 488/405 nm excitation ratio was used to calculate ATP/ADP ratios. pHrodo-Red was excited at 561 nm with emission collected at 585 nm. Following 2 minutes of baseline recording, treatments were applied using an electronic pipette (Eppendorf Repeater E3). Neurons were then imaged for 6 minutes in the presence of drug-containing HL-3. Following a 2-minute drug washout period, 10 mM NH_4_Cl was applied for empirical pH calibration as described (30). For each experimental condition, 15-20 neurons were imaged per dish across a minimum of 3 biological replicates.

#### Image quantification with pH correction

For each field of view, we obtained fluorescence emissions from ROIs corresponding to individual neuronal cell bodies. To correct for background fluorescence, we subtracted the intensities of an ROI lacking cells from the intensities of the neuronal ROIs. Next, we corrected the emission intensities for fluorescence bleach, and then calculated the PercevalHR ratios with the bleach-corrected intensities. We normalized the ratios to the means of their corresponding baseline values. We utilized an empirical pH correction protocol (30) that first normalized the PercevalHR ratios to the exponential fits of the premixing baselines in order to set the average premixing baselines to 1. NH_4_Cl-induced deviations of the pHrodo and PercevalHR ratio signals from baseline exhibited linear relationships, which were used to empirically correct for the changes in sensor ratios that stem solely from changes in cytosolic pH. We determined the integrated changes in normalized, pH-corrected ratios by calculating the area under the curve (AUC) for each trace from the time of treatment to washout. We used the CausalImpact package (56) in R to extrapolate the baseline, and thereby, estimated the ratios had we not applied the treatment. Difference between the AUC value of a trace and that of its extrapolated baseline, adjusted for the length of time after the treatment, represented the integrated change in ratios.

### Seahorse metabolic flux analyses and quantification

#### Extracellular flux analysis

Metabolic flux analyses were conducted on *ex vivo* 3^rd^ instar larval brains using a Seahorse XF HS Mini analyzer (Agilent Technologies) with XF HS Mini cell culture plates (Agilent) and XFp sensor cartridges (Agilent). Sensor cartridges were hydrated overnight in sterile water at 25°C in a non-CO_2_ incubator. The following day, hydration solution was replaced with XF Calibrant (Agilent) and cartridges were incubated for an additional 24 hours at 25°C.

#### Assay medium and drug preparation

Assay medium was phenol red-free XF DMEM (Agilent) supplemented with either 10 mM glucose alone or 10 mM glucose, 1 mM sodium pyruvate and 2 mM glutamine (full). Medium pH was adjusted to 7.4 with NaOH. Drug stocks were diluted in the assay media to 30 μM oligomycin A (Cayman Chemicals), 3 μM FCCP (Sigma Aldrich), 10 μM rotenone (Sigma Aldrich) and 10 μM antimycin A (Sigma Aldrich) and 200 mM 2DG (Thermo Scientific).

#### Tissue preparation

During sensor cartridge calibration, 3^rd^ instar larval brains were dissected in the assay medium and transferred to XFp cell culture plates pre-coated with Cell-Tak cell and tissue adhesive (Corning). Individual brains were placed in wells containing 50 μl assay medium. Custom circular tissue restraints fabricated from nylon mesh (Instrument Design and Fabrication Core, Arizona State University) were positioned over each brain to prevent tissue displacement during measurements. Plates were equilibrated at 25°C for 15 minutes before initiating recordings.

#### Data acquisition

All measurements were performed at 25°C. Basal oxygen consumption rate (OCR) and extracellular acidification rate (ECAR) were measured at baseline and following sequential injection of drugs. Data were extracted using Wave Desktop Software (Agilent).

#### Analysis of ECAR data

ECAR data from Seahorse experiments were analyzed using linear mixed-effects models in R using the lme4 package (57). Data from individual wells were grouped by genotype and experimental batch to account for variation between runs. For analysis of glycolytic reserve capacity, ECAR measurements from baseline and rotenone + antimycin A treatment phases were modeled using phase (coded 0 for baseline and 1 for post-treatment) as a fixed effect with genotype interaction terms. The model included random intercepts for individual wells and experimental batch to account for repeated measures and batch effects: ECAR ∼ phase × genotype + (1|well) + (1|batch). Maximal glycolytic capacity was assessed separately by modeling ECAR values during the rotenone + antimycin A phase, using genotype as a fixed effect with well and batch as random intercepts to account for repeated measures and experimental batch effects: ECAR ∼ genotype + (1|well) + (1|batch). Estimated marginal means from this model were used to test for genotype differences in maximal ECAR.

To assess the response to glycolytic inhibition, ECAR decline following 2-DG injection was analyzed using time-centered values with genotype as an interaction term. The mixed-effects model incorporated random intercepts for wells and batch: ECAR ∼ time × genotype + (1|well) + (1|batch). Estimated marginal means and slopes were calculated using the emmeans package to determine group-specific responses. For glycolytic reserve, pairwise contrasts between phases within each genotype were computed. For 2-DG response, the rate of ECAR decline (slope) was compared between genotypes.

#### Analysis of OCR data

ATP-linked respiration was assessed by comparing Seahorse OCR values before and after oligomycin A treatment. The mixed-effects model structure was: OCR ∼ phase × genotype + (1|well) + (1|batch), where phase was coded as 0 (baseline) or 1 (post-oligomycin). ATP-linked respiration was calculated as the difference between baseline and post-oligomycin OCR for each genotype.

Respiratory reserve was determined by comparing baseline OCR values and OCR after FCCP uncoupling. The model used phase coding of 0 (baseline) and 1 (FCCP). Maximal respiratory capacity was determined by comparing OCR values after FCCP uncoupling versus non-mitochondrial respiration (post-rotenone + antimycin A). The model used phase coding of 0 (rotenone + antimycin A) and 1 (FCCP). Individual wells were grouped by genotype and assigned to experimental batches. Random intercepts for wells and batch controlled for repeated measures and technical variation between runs. Estimated marginal means were calculated for each genotype at each treatment phase. Pairwise contrasts within genotypes determined the magnitude of oligomycin-sensitive (ATP-linked) respiration, respiratory reserve and maximal respiratory capacity. Between-group differences were then assessed.

### Lifespan analysis

Newly eclosed adult flies were collected and transferred to vials containing standard fly food at a density of ≤ 15 flies per vial. Flies were maintained at room temperature and transferred to new vials containing fresh fly food twice weekly. During each transfer, the number of dead flies were counted until all the animals in that cohort had died. Kaplan-Meier survival plots were generated using GraphPad Prism.

#### Gompertz mortality modeling

Empirical hazard functions were estimated from survival data using the muhaz package in R with global bandwidth selection. For each genotype, log-transformed hazard rates were fitted to a linear model using ordinary least squares regression. Model fit was assessed using R^2^ values. Age-specific mortality was then modeled using the Gompertz distribution with the flexsurv package (58) in R. Maximum likelihood estimation was performed with the Gompertz distribution for each genotype separately. Two key parameters were extracted, the aging rate and the baseline mortality at the start of adult life. Between-genotype differences in Gompertz parameters were assessed using z-tests. For each parameter comparison, the test statistic was calculated as z = (parameter_1_ – parameter_2_) / (SE_1_^2^ + SE_2_^2^)^1/2^, with standard errors obtained from the maximum likelihood estimation. Two-tailed *P*-values were calculated from the standard normal distribution. All pairwise comparisons between genotypes were performed, with *P*-value adjustments for multiple testing using the Holm-Bonferroni method.

---

**Figure S1. Overexpression of *itpr* produces transcriptional changes overlapping with the tau^0N3R^ signature.**

**(A)** Volcano plots showing DEGs in larval brains overexpressing *itpr* (*tubulin>itpr*) relative to *UAS-itpr*/+ (left) and *tubulin-GAL4*/+ (right) controls. Upregulated genes (teal), downregulated genes (magenta), and non-significant genes (gray, adjusted *P*-value ≥ 0.05) are indicated. Also shown is the *itpr* gene (red), which is significantly upregulated.

**(B)** Scatter plot comparing log_2_ fold changes of DEGs identified relative to each control line. High concordance (slope ∼ 1) and 98.6% directional consistency demonstrate robust transcriptional changes induced by *itpr* overexpression.

**(C)** Venn diagram showing overlap between tau^0N3R^ DEGs (green, 4,519 genes) and *itpr* overexpression DEGs (blue, 627 genes). 475 genes (76% of itpr DEGs) are shared, representing a highly significant enrichment.

**(D)** Scatter plot directly comparing log_2_ fold changes between DEGs in *tau^0N3R^* and *itpr* overexpression (o/e) for all genes.

**(E)** Gene Ontology enrichment analysis of the 475 shared downregulated genes. Bar length represents fold enrichment, and color intensity indicates FDR.

**Figure S2. Glycolytic gene knockdown in *tau^0N3R^*-expressing animals.**

**(A and C)** Kaplan-Meier survival curves in the indicated genotypes. Dotted line indicates 50% survival. See Supplemental Table 8A for sample sizes.

**(B)** Quantitative RT-PCR validation of RNAi-mediated transcript. All values are normalized to *GAL4*/+ controls and represent mean ± SEM, n = 3-6 biological replicates.

\**P* < 0.05, \*\**P* < 0.01, \*\*\**P* < 0.001, \*\*\*\**P* < 0.0001, one-sample t-tests against control mean (1.0).

**Figure S3. Glycolytic gene knockdown in control neurons does not alter aging kinetics.**

**(A-B)** Gompertz aging rate parameter **(A)** and baseline mortalities **(B)** in the indicated genotypes. Values represent mean ± standard error from Gompertz model fits on lifespan data. All statistical comparisons were made with *d42>tau^0N3R^* flies. \*\*\*\**P* < 0.0001, \*\**P* < 0.01, n.s., not significant, Wald z-test on pairwise parameter differences with Holm post hoc correction for multiple comparisons.

## Notes

### Competing Interest Statement

The authors have declared no competing interest.

## REFERENCES

1. P. Lei, S. Ayton, A. I. Bush, The essential elements of Alzheimer’s disease. J Biol Chem 296, 100105 (2021).

2. V. Crowell, et al., Disease severity and mortality in Alzheimer’s disease: an analysis using the U.S. National Alzheimer’s Coordinating Center Uniform Data Set. BMC Neurol 23, 302 (2023).

3. C.-S. Liang, et al., Mortality rates in Alzheimer’s disease and non-Alzheimer’s dementias: a systematic review and meta-analysis. Lancet Healthy Longev 2, e479–e488 (2021).

4. S. Wang, et al., Spatiotemporal transcriptomic profiling reveals metabolic dysfunction prior to overt tauopathy in the PS19 mouse model. Res Sq (2025). 10.21203/rs.3.rs-6941464/v1.

5. A. Pichet Binette, et al., Proteomic changes in Alzheimer’s disease associated with progressive Aβ plaque and tau tangle pathologies. Nat Neurosci 27, 1880– 1891 (2024).

6. S. M. de la Monte, M. Tong, Brain metabolic dysfunction at the core of Alzheimer’s disease. Biochem Pharmacol 88, 548–59 (2014).

7. F. Marinaro, et al., Molecular and cellular pathology of monogenic Alzheimer’s disease at single cell resolution. bioRxiv [Preprint] (2020). Available at: http://biorxiv.org/lookup/doi/10.1101/2020.07.14.202317 [Accessed 13 October 2025].

8. L. Traxler, et al., Warburg-like metabolic transformation underlies neuronal degeneration in sporadic Alzheimer’s disease. Cell Metab 34, 1248–1263.e6 (2022).

9. J. Mertens, et al., Age-dependent instability of mature neuronal fate in induced neurons from Alzheimer’s patients. Cell Stem Cell 28, 1533–1548.e6 (2021).

10. S. Garcia, et al., Enhanced glycolysis and GSK3 inactivation promote brain metabolic adaptations following neuronal mitochondrial stress. Hum Mol Genet 31, 692–704 (2022).

11. M. C. B. D’Alessandro, S. Kanaan, M. Geller, D. Praticò, J. P. L. Daher, Mitochondrial dysfunction in Alzheimer’s disease. Ageing Res Rev 107, 102713 (2025).

12. L. Traxler, et al., Metabolism navigates neural cell fate in development, aging and neurodegeneration. Dis Model Mech 14, dmm048993 (2021).

13. C.-O. Wong, et al., Regulation of longevity by depolarization-induced activation of PLC-β-IP3R signaling in neurons. Proc Natl Acad Sci U S A 118 (2021).

14. S. D. Pletcher, A. A. Khazaeli, J. W. Curtsinger, Why Do Life Spans Differ? Partitioning Mean Longevity Differences in Terms of Age-Specific Mortality Parameters. J Gerontol A Biol Sci Med Sci 55, B381–B389 (2000).

15. L. D. Mueller, T. J. Nusbaum, M. R. Rose, The Gompertz equation as a predictive tool in demography. Exp Gerontol 30, 553–569 (1995).

16. W. Mair, P. Goymer, S. D. Pletcher, L. Partridge, Demography of Dietary Restriction and Death in *Drosophila*. Science (1979) 301, 1731–1733 (2003).

17. A. Mudher, et al., GSK-3beta inhibition reverses axonal transport defects and behavioural phenotypes in Drosophila. Mol Psychiatry 9, 522–30 (2004).

18. J. Piñero, et al., The DisGeNET knowledge platform for disease genomics: 2019 update. Nucleic Acids Res 48, D845–D855 (2020).

19. J. Piñero, et al., DisGeNET: a comprehensive platform integrating information on human disease-associated genes and variants. Nucleic Acids Res 45, D833– D839 (2017).

20. Y. Hu, et al., An integrative approach to ortholog prediction for disease-focused and other functional studies. BMC Bioinformatics 12, 357 (2011).

21. A. Becker, P. Schlöder, J. E. Steele, G. Wegener, The regulation of trehalose metabolism in insects. Experientia 52, 433–9 (1996).

22. G. R. Wyatt, G. F. Kale, The chemistry of insect hemolymph. II. Trehalose and other carbohydrates. J Gen Physiol 40, 833–47 (1957).

23. T. Yasugi, T. Yamada, T. Nishimura, Adaptation to dietary conditions by trehalose metabolism in Drosophila. Sci Rep 7, 1619 (2017).

24. T. TeSlaa, M. Ralser, J. Fan, J. D. Rabinowitz, The pentose phosphate pathway in health and disease. Nat Metab 5, 1275–1289 (2023).

25. J. Yip, X. Geng, J. Shen, Y. Ding, Cerebral Gluconeogenesis and Diseases. Front Pharmacol 7, 521 (2016).

26. S. Bawa, et al., Drosophila TRIM32 cooperates with glycolytic enzymes to promote cell growth. Elife 9 (2020).

27. N. Raun, et al., Trithorax regulates long-term memory in Drosophila through epigenetic maintenance of mushroom body metabolic state and translation capacity. PLoS Biol 23, e3003004 (2025).

28. S. Spannl, et al., Glycolysis regulates Hedgehog signalling via the plasma membrane potential. EMBO J 39, e101767 (2020).

29. N. E. Karagas, et al., Loss of Activity-Induced Mitochondrial ATP Production Underlies the Synaptic Defects in a Drosophila Model of ALS. J Neurosci 42, 8019–8037 (2022).

30. M. S. Price, et al., Intracellular lactate dynamics in Drosophila neurons. iScience 28, 113462 (2025).

31. O. Warburg, On the origin of cancer cells. Science 123, 309–14 (1956).

32. J. Yao, et al., Human tau promotes Warburg effect-like glycolytic metabolism under acute hyperglycemia conditions. J Biol Chem 301, 108376 (2025).

33. K. A. Hughes, B. Charlesworth, A genetic analysis of senescence in Drosophila. Nature 367, 64–6 (1994).

34. R. M. Denton, Regulation of mitochondrial dehydrogenases by calcium ions. Biochim Biophys Acta Bioenerg 1787, 1309–1316 (2009).

35. M. E. Levine, et al., An epigenetic biomarker of aging for lifespan and healthspan. Aging 10, 573–591 (2018).

36. M. E. Levine, A. T. Lu, D. A. Bennett, S. Horvath, Epigenetic age of the pre-frontal cortex is associated with neuritic plaques, amyloid load, and Alzheimer’s disease related cognitive functioning. Aging 7, 1198–1211 (2015).

37. C. Gaser, K. Franke, S. Klöppel, N. Koutsouleris, H. Sauer, BrainAGE in Mild Cognitive Impaired Patients: Predicting the Conversion to Alzheimer’s Disease. PLoS One 8, e67346 (2013).

38. W. Zhou, et al., Neuronal aerobic glycolysis exacerbates synapse loss in aging mice. Exp Neurol 371, 114590 (2024).

39. J. Yang, et al., Lactate promotes plasticity gene expression by potentiating NMDA signaling in neurons. Proc Natl Acad Sci U S A 111, 12228–12233 (2014).

40. D. Zhang, et al., Metabolic regulation of gene expression by histone lactylation. Nature 574, 575–580 (2019).

41. R.-Y. Pan, et al., Positive feedback regulation of microglial glucose metabolism by histone H4 lysine 12 lactylation in Alzheimer’s disease. Cell Metab 34, 634–648.e6 (2022).

42. D. M. Long, et al., Lactate dehydrogenase expression modulates longevity and neurodegeneration in Drosophila melanogaster. Aging 12, 10041–10058 (2020).

43. D. Jimenez-Blasco, R. Lapresa, J. Agulla, A. Almeida, J. P. Bolaños, Neuronal glycolysis meets mitophagy to govern organismal wellbeing. Trends in Endocrinology & Metabolism (2025). 10.1016/J.TEM.2025.05.005.

44. D. L. Croteau, E. F. Fang, H. Nilsen, V. A. Bohr, NAD+ in DNA repair and mitochondrial maintenance. Cell Cycle 16, 491–492 (2017).

45. A. Herrero-Mendez, et al., The bioenergetic and antioxidant status of neurons is controlled by continuous degradation of a key glycolytic enzyme by APC/C–Cdh1. Nat Cell Biol 11, 747–752 (2009).

46. Y.-J. Park, et al., Distinct systemic impacts of Aβ42 and Tau revealed by whole-organism snRNA-seq. Neuron 113, 2065–2082.e8 (2025).

47. L. Liu, K. R. MacKenzie, N. Putluri, M. Maletić-Savatić, H. J. Bellen, The Glia-Neuron Lactate Shuttle and Elevated ROS Promote Lipid Synthesis in Neurons and Lipid Droplet Accumulation in Glia via APOE/D. Cell Metab 26, 719–737.e6 (2017).

48. A. Volkenhoff, et al., Glial Glycolysis Is Essential for Neuronal Survival in Drosophila. Cell Metab 22, 437–447 (2015).

49. T. L. Parkes, et al., Extension of Drosophila lifespan by overexpression of human SOD1 in motorneurons. Nat Genet 19, 171–174 (1998).

50. T. Lee, L. Luo, Mosaic analysis with a repressible cell marker for studies of gene function in neuronal morphogenesis. Neuron 22, 451–61 (1999).

51. K. Tenney, et al., Drosophila Rtf1 functions in histone methylation, gene expression, and Notch signaling. Proc Natl Acad Sci U S A 103, 11970–11974 (2006).

52. K. Venkatesh, G. Siddhartha, R. Joshi, S. Patel, G. Hasan, Interactions between the inositol 1,4,5-trisphosphate and cyclic AMP signaling pathways regulate larval molting in Drosophila. Genetics 158, 309–18 (2001).

53. Y. Liao, G. K. Smyth, W. Shi, The R package Rsubread is easier, faster, cheaper and better for alignment and quantification of RNA sequencing reads. Nucleic Acids Res 47, e47–e47 (2019).

54. Y. Tang, M. Horikoshi, W. Li, ggfortify: Unified Interface to Visualize Statistical Results of Popular R Packages. R J 8, 474 (2016).

55. M. I. Love, W. Huber, S. Anders, Moderated estimation of fold change and dispersion for RNA-seq data with DESeq2. Genome Biol 15, 550 (2014).

56. K. Brodersen, F. Gallusser, J. Koehler, N. Remy, Inferring causal impact using Bayesian structural time-series models. (2015).

57. D. Bates, M. Mächler, B. Bolker, S. Walker, Fitting Linear Mixed-Effects Models Using **lme4**. J Stat Softw 67, 1–48 (2015).

58. C. Jackson, flexsurv : A Platform for Parametric Survival Modeling in *R*. J Stat Softw 70, 1–33 (2016).

