## Supplementary figures and images for "Neuronal glycolytic reprogramming drives lethality via accelerated aging in a *Drosophila* model of tauopathy"

### Supplemental Figure S1

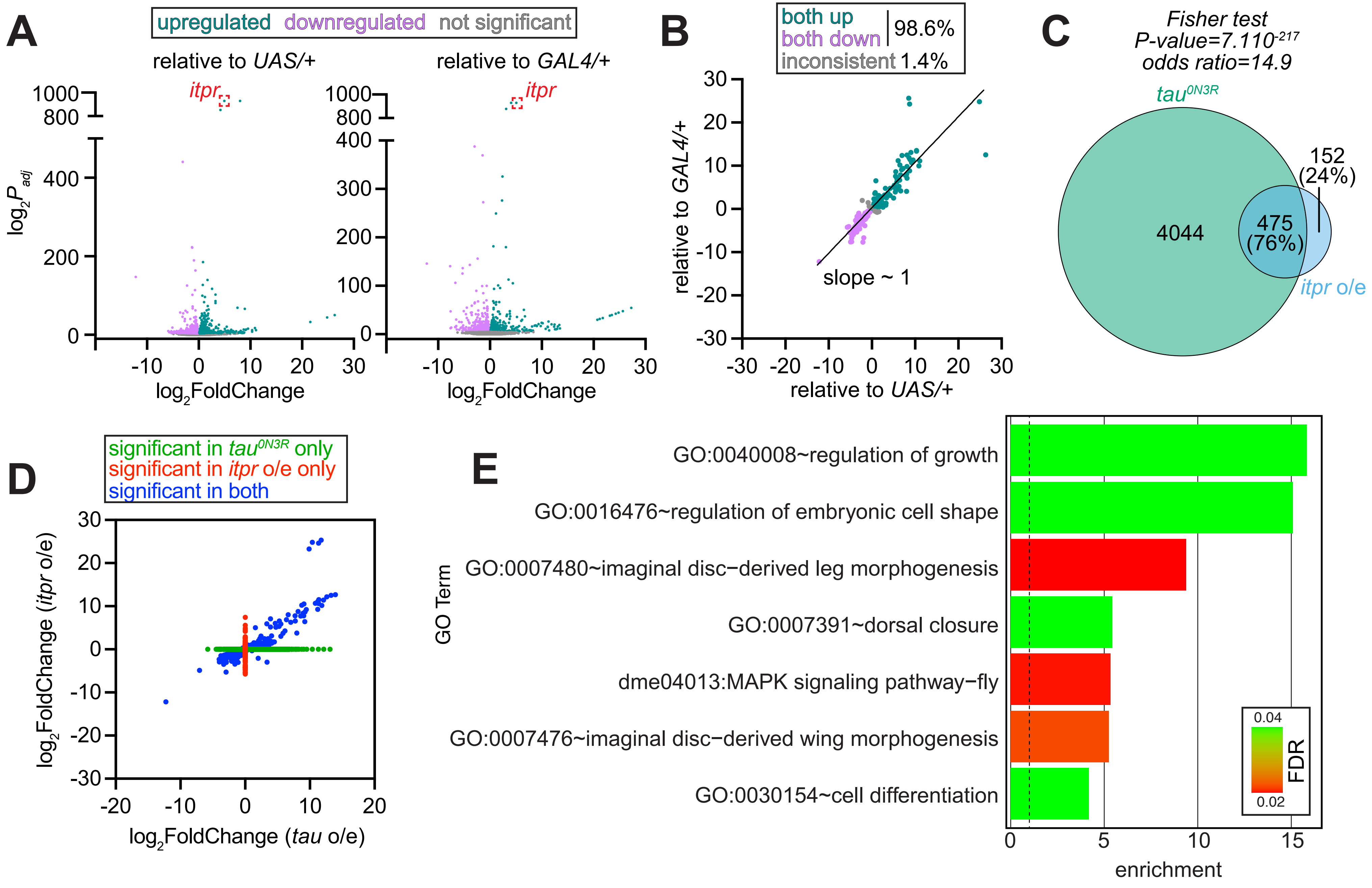

### Supplemental Figure S2

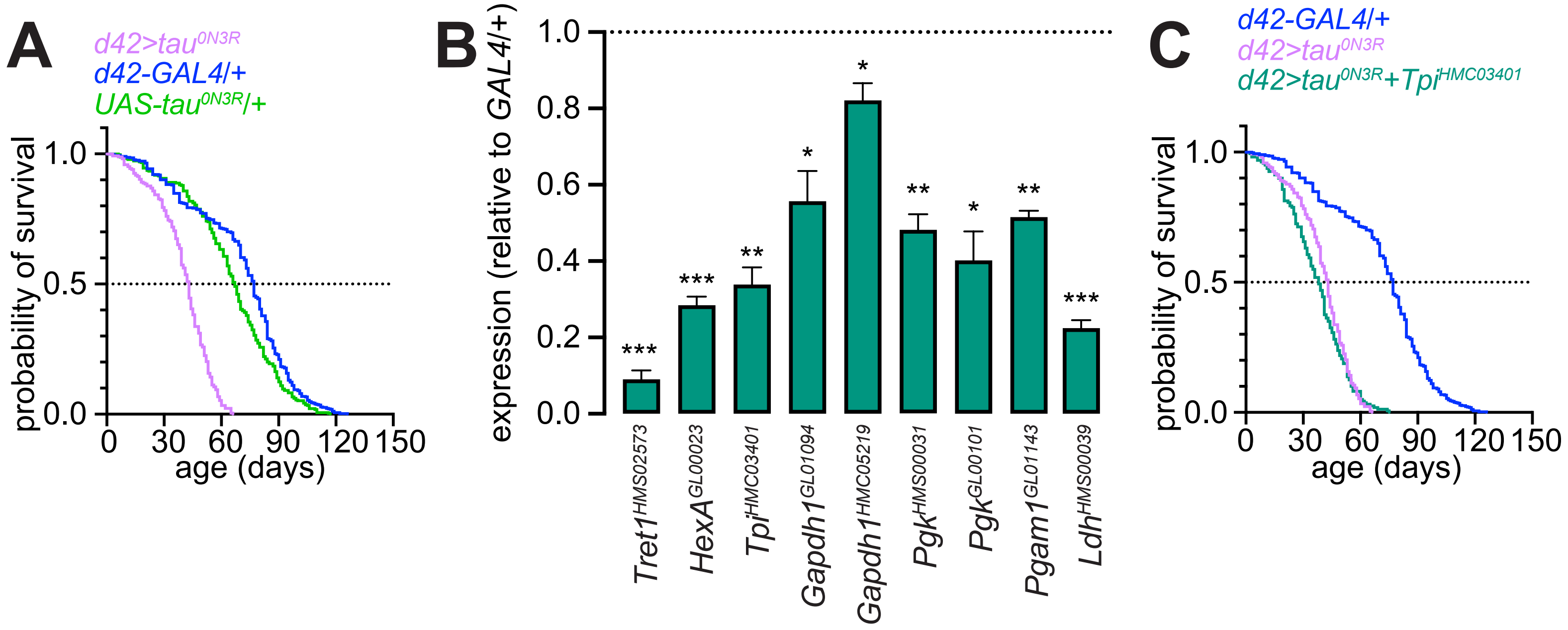

### Supplemental Figure S3

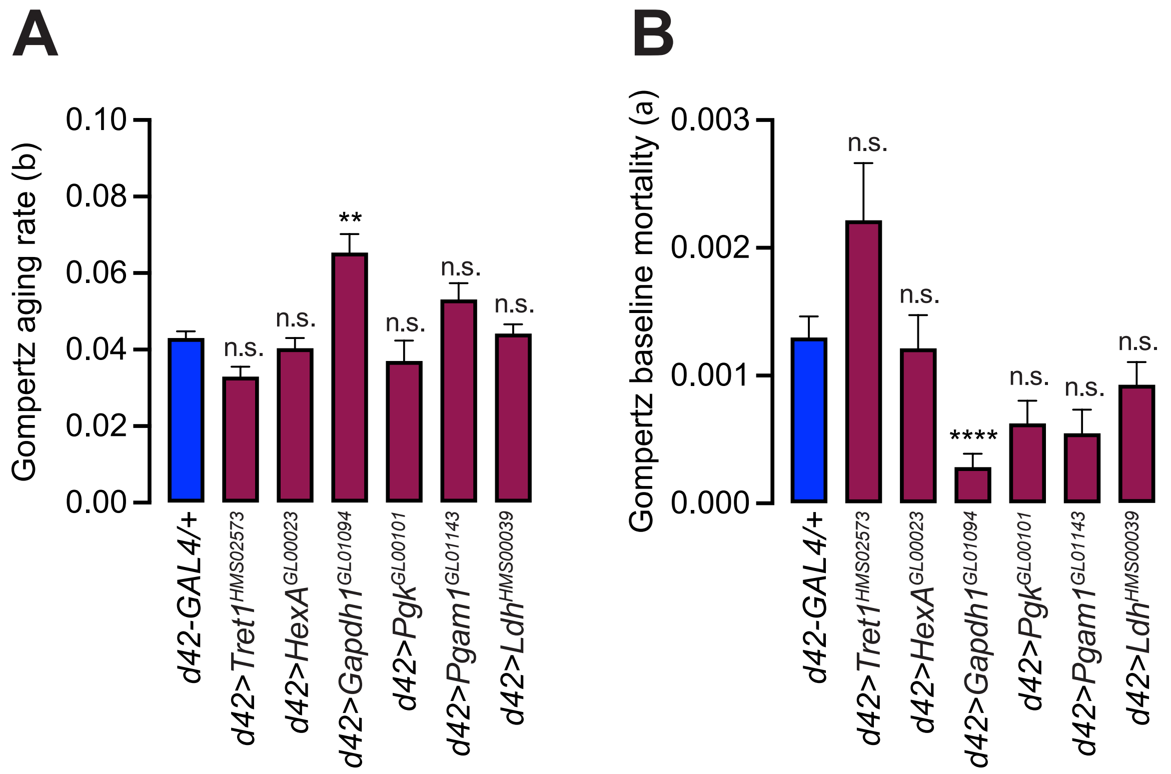
